# Using a Deep Learning Approach for Model-based Control of Deep Brain Stimulation

**DOI:** 10.1101/2024.10.29.620970

**Authors:** Srikar Saradhi, Yupeng Tian, Mohammed R. Rezaei, Milad Lankarany

## Abstract

Deep brain stimulation (DBS) has been developed as a treatment method for various neurological disorders, including Parkinson’s disease, essential tremor and depression. Although DBS is effective, it often loses efficacy over sustained periods because a constant stimulation is applied without adapting to the patient’s current clinical state. In contrast, an adaptive closed-loop DBS system can offer more tailored stimulation in real-time based on a feedback biomarker. In early 2024, we developed a model-based DBS control framework that consists of three main functions: (1) a biophysically reasonable encoding model, (2) a simple decoding model, and (3) a controller. We used a polynomial fit function in the decoding model to approximate the neural-motor relationship, from DBS-induced Vim neural activity to muscle fiber electromyography (EMG). Despite promising results, the polynomial method is inaccurate in capturing the full representation of the neural-motor (EMG) function across different DBS frequencies. In this work, to capture the nonlinear intricate relationship between the neural and EMG patterns, we developed a one-dimensional convolutional neural network (1-D CNN) as a decoding model to predict the EMG signal directly from the DBS-induced Vim neural activity. The 1-D CNN network outputted a high R^2^ value of 0.997 which significantly outperformed the polynomial method (R^2^ = 0.277) and a deep learning approach based on long short-term memory (R^2^ = 0.296). We anticipate that our work highlights the need for a data-driven approach that can reliably map neural activities to symptomatic signals like EMG for better adjusting DBS parameters.

## Introduction

Deep brain stimulation (DBS) has been developed as a treatment method for various neurological disorders, including Parkinson’s disease [1], essential tremor [2] and depression [3]. DBS of the thalamic ventral intermediate nucleus (Vim) has been a standard therapy for essential tremor (ET) [4][2][5]. In current clinical practice, DBS is manually programmed and delivered to the patient in an open-loop continuous mode, relying solely on the expertise and judgment of neurologists [6][7][8]. Such continuous DBS methods (cDBS) have been associated with potential side effects, increased stimulation habituation, and reduced battery longevity [9][10][11][12][13]. Thus, there is an imperative need for a closed-loop DBS system delivering on-demand stimulations in response to essential tremor symptoms [14][15][9].

However, closed-loop DBS requires a biomarker that reflects the patient’s current clinical state. This biomarker can be used to tune the stimulation frequency, intensity, and pulse width of the DBS system [14] [15][16][17][18]. For instance, the modulation of local field potential (LFP) beta band power within a specific range is often employed as a control mechanism for Parkinson’s disease [17]. However, such approaches are hindered by a limited understanding of the underlying physiological mechanisms of DBS and the neuronal circuits implicated in disease pathology [8][16]. Thus, it is imperative to develop methodological frameworks to link DBS-modulated neural activity to symptomatic behaviors [16][19][20].

In our recent work (Tian et al., 2024), we developed a data-driven DBS framework that traces the neural-motor pathway from DBS-induced Vim neural activity to muscle fiber electromyography (EMG). Short-term synaptic plasticity was captured by following spike trains from M1 cortical neurons through spinal motor neurons to the muscle fibers. The direct simulation of EMG with the encoding computational model is time-consuming and impractical for a biomarker-based adaptive controller. A polynomial method was used to decode the EMG signal directly from the DBS-induced Vim firing rates to enable a quasi-real-time computation of the EMG biomarker for control applications [16]. However, the closed-loop system in Tian et al. (2024) [16] left several key problems unsolved: (1) The simulated EMG misses some critical features observed in clinical EMG data; (2) The polynomial method is inadequate in accurately decoding the EMG biomarker, highlighting the need for a robust data-driven approach that can reliably predict symptomatic signals.

In this work, we develop a one-dimensional convolutional neural network (1-D CNN) as a decoding model to predict the EMG biomarker directly from the DBS-induced Vim neural activity, facilitating the potential implementation of a closed-loop control system [16].

The rationale behind investigating deep learning is its ability to capture the nonlinear intricate relationship between the neural and EMG patterns [21][22] for different DBS frequencies. The challenges of deep learning approaches are twofold, one being that its performance and decision-making are often limited in explainability [23], and the other being that it requires a large dataset for effective training [24]. The first issue can be addressed by keeping the model architecture as simple as possible by ensuring that each hidden layer in the deep learning model has a clear, interpretable function [23]. The second issue is addressed by generating a large, high-quality dataset using our encoding model, developed in our previous work, to simulate DBS-induced Vim neural activity and corresponding EMG data. Our 1-D CNN model can capture the relationship between Vim spiking and EMG output for different DBS frequencies (20 – 200 Hz). The 1-D CNN-decoded EMG biomarker could be potentially implemented in a closed-loop controller (e.g., proportional-integral-derivative (PID) [25][26][27]) that automatically updates the appropriate DBS frequency. The data-driven deep learning decoding model developed in this work is feasible for an adaptive controller because of its higher accuracy and efficacy in comparison to the polynomial-based decoding model developed in our recent work (Tian et al., 2024). The power spectrum should be computed directly from the EMG as a biomarker to be effectively used for an adaptive controller. Unlike the polynomial fit method, the deep learning model, once trained, can directly approximate this from the processed spectrogram features. The impact of this study is that the developed framework can be implemented in and out of the clinic to continuously adapt and adjust the appropriate patient-specific DBS frequencies to effectively suppress essential tremor symptoms.

## Methods

The data-driven DBS system consists of two key models: (1) The encoding computational model developed by Tian et al. (2024) which computes Vim spike trains from DBS and generates simulations of clinically observed EMG features such as voluntary muscle activities [28][9] and bipolar symmetry [29][30]; (2) The data-driven deep learning model to predict the EMG biomarker by tracking the simulated EMG provided by the encoding model. The decoded EMG biomarker could then be used in a closed-loop controller to automatically update the appropriate DBS frequency [16]. **Figure 1** illustrates the schematics of the model-based DBS system.

**Figure 1.**
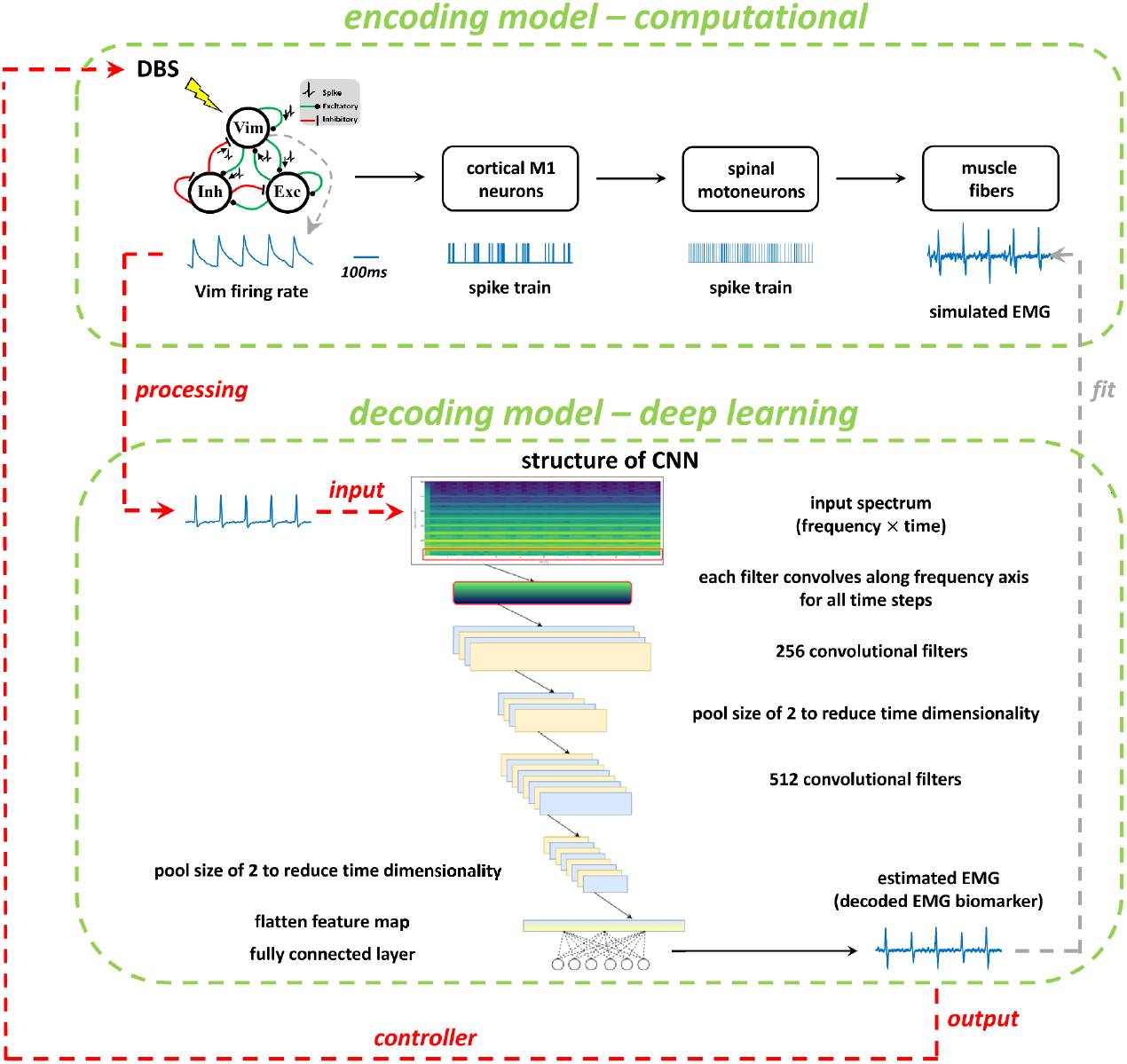
Schematic illustration of the model-based DBS control framework. The controller outputs a frequency of DBS pulses, which are used to obtain the Vim firing rate from the computational encoding model. The Vim firing rate is first processed, then inputted into a 1-D CNN model to decode an estimated EMG signal. The estimated EMG signal is the output biomarker, which is processed by a closed-loop controller to find the appropriate DBS frequency [16]. The CNN structure diagram displays the flow of information from the input (represented as a spectrogram) to its processed feature maps using an initial layer of 256 filters with a kernel size of 7 followed by a deeper layer of 512 filters with a kernel size of 3, both with a stride of 1. Finally, the feature maps are processed through a fully connected layer which produces an output EMG signal.

### Encoding Model – A Biophysically Reasonable Model

The encoding model consists of four parts as shown in **Figure 1**: (*i*) the Vim-network model of the effects of Vim-DBS; (*ii*) motor cortical neuron spikes influenced by DBS and background activities; (*iii*) spinal motoneurons activities impacted by neurons in the motor cortex; (*iv*) motor unit action potentials in the muscle fibers innervated by the spinal motoneurons.

DBS-impacted Vim firing rate was simulated by our previous Vim-network model that accurately reproduced the clinically recorded instantaneous firing rate of the Vim neurons receiving DBS of stimulation frequencies ranging from 10 to 200 Hz [31]. During Vim-DBS, the activities of a neuron in the primary motor cortex (M1) are modeled as being influenced by two factors: (1) the direct activation of the Vim-M1 axon by Vim-DBS; and (2) the background activities that induce the tremor symptom and voluntary muscle movements. Vim-DBS activates the Vim-M1 axons connecting to the synapses projecting to the M1 neuron, and these synapses are characterized by the Tsodyks & Markram (TM) model [32]. Besides the direct Vim-M1 axon activation, the M1 neuron is also affected by background activities, including the spikes that are (a) from the DBS-induced firings of the Vim neurons, (b) inducing tremor symptoms observed when DBS-OFF, and (c) generating voluntary muscle movements. Because essential tremor is often in the frequency band 4 – 8 Hz [33][34][18], the tremor-inducing background firing rate was taken as a waveform consisting of a baseline shift and 6-Hz bursts. Each burst consists of 3 consecutive 20-ms sinusoidal waves, to be consistent with observed tremors in clinical EMG recordings [33][35][34]. The voluntary muscle movements were modeled by a sinusoidal firing rate waveform with a mean rate of 5 Hz [28][36], and oscillation frequency at 0.5 Hz, to fit the voluntary muscle oscillations observed in the EMG recording from essential tremor patient in Cenera et al. (2021) [9]. The total background firing rate waveform is the sum of the firing rate from the above (a), (b), and (c). Then, this total waveform is used as the time-varying Poisson rate for generating Poisson spike trains as background inputs to an M1 neuron, whose membrane potential was characterized by a leaky integrate-and-fire (LIF) model:

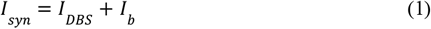

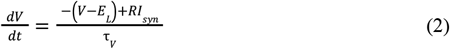

where *E*_*L*_= −65 mV is the equilibrium potential, *R* (resistance parameter) = 1 MΩ, and τ_*V*_ = 10 ms is the membrane time constant; spikes occur when *V* ≥ *V*_*th*_, where *V*_*th*_ = −35 mV. The reset voltage is −90 mV and the absolute refractory period is 1 ms. *I*_*DBS*_ is the input induced by direct Vim-M1 axon activation by DBS, and *I*_*b*_ is the input from background spikes. *I*_*DBS*_ and *I*_*b*_ are obtained by characterizing the M1 synapse with the TM model [32]. *I*_*syn*_ is the total post-synaptic input current (**Equation (1)**). Our model of the Vim-DBS effects on the cortical M1 neurons was validated by comparing with 10-s of neural spike data from non-human primate single-unit recordings in M1 during 130-Hz VPLo-DBS (equivalent to Vim-DBS in humans [37]), reported in Agnesi et al. (2015) [38][16].

The spikes of M1 neurons induce activities in the motor neurons in the spinal cord [36]. The membrane potential of the spinal motor neuron is given by a LIF model equivalent to that of Herrmann and Gerstner (2002) [39][36]:

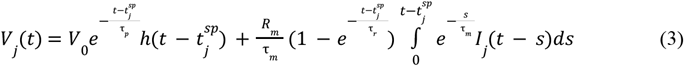

where 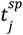 is the last spike timing of the j^th^ motor neuron. *h*(*t*) is the Heaviside step function, which is zero when its argument is negative and one otherwise. *I*_*j*_ (*t*) is the postsynaptic current from M1 neurons into the j^th^ motor neuron. *V*_0_ =− 22mV is the resting membrane potential [36]. *R*_*m*_ = 36 MΩ is the input resistance [36]. τ_*p*_ = 2 ms is the refractory time constant, τ_*m*_ = 4 ms is the passive membrane time constant, τ_*r*_ = 100 ms is the recovery time constant [40][36]. The firing of motor neurons is determined by an exponential distribution of firing thresholds (**Equations (4) and (5)**) [28]:

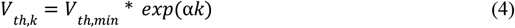

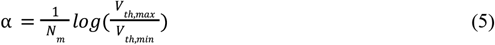

*V*_*th,k*_ is the firing threshold of the *k*^th^ motor neuron. When the membrane potential of a motor neuron reaches a firing threshold, it is instantaneously reset to resting potential *V*_0_ (**Equation (3)**), and the integration process restarts. *V*_*th,min*_ = 10 mV is the minimal firing threshold, and *V*_*th,max*_ = 900 mV is the maximal firing threshold [28]. α is a scaling parameter, and *N*_*m*_ =120 is the number of motor neurons.

The spikes from the spinal motor neurons generate action potentials in the motor units of muscle fibers [28][36][41]. The shape of motor unit action potential (MUAP) was characterized by a first-order Hermite-Rodriguez function [42][28][36]:

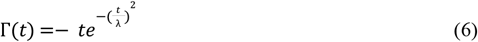

where λ = 2 ms is a constant timescale [42][36]. For each motor neuron, the scaling amplitude of the corresponding MUAP is proportional to the firing threshold of the motor neuron [28]. The MUAP is modeled as:

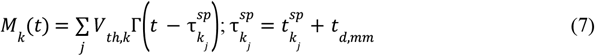

where *M*_*k*_ is the MUAP of the *k*^th^ motor unit, corresponding to the *k*^th^ motor neuron. *V*_*th,k*_ is the firing threshold of the *k*^th^ motor neuron. 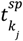 is the *j*^th^ spike timing of the *k*^th^ motor neuron, and *t*_*d,mm*_ = 10 ms is the motor neuron-to-muscle conduction delay in humans [43]. The EMG time series was modeled as the sum of MUAP innervated by each motor neuron:

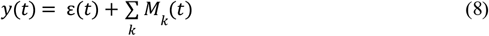

where *y*(*t*) is the modeled EMG, and ε(*t*) is a low-level Gaussian white noise [41][36] with a standard deviation of 1/300 mV.

### Data-Driven Model – A Deep Learning Model

The 1-D CNN model was trained and evaluated on the simulated Vim firing rate and EMG data from the encoding model for various DBS frequencies. The input to the deep learning model is a one-step processing of the simulated Vim firing rate (**Figure 1**) with the following convolution:

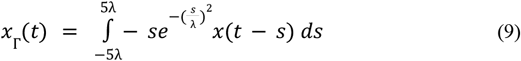

x is the simulated Vim firing rate from the encoding model, x_Γ is the processed signal by convolving with the MUAP shape kernel, Γ (**Equation (6)**), λ = 2 ms is a constant timescale [42][36]. The 1-D CNN model is fitted to the output reference data, which is the simulated EMG signals (**Equation (8), Figure 1**).

Ten seconds of data were simulated for DBS frequencies ranging from 20 Hz to 200 Hz in 10 Hz increments, using a sampling frequency of 10,000 Hz, resulting in a total of 20 input-output simulations. This range of DBS frequencies was selected to ensure robustness in the model’s ability to differentiate between various stimulation frequencies that are commonly used in DBS treatment [44]. This simulation process was repeated five times with each iteration incorporating a different randomness noise distribution to introduce variability. Data from three of the simulations were concatenated and used for training [45]. The fourth simulation was used for validation and the fifth simulation was used for testing. The testing data covers the same range of DBS frequencies but in 5 Hz incremental steps to further assess the model’s ability to capture the underlying relationship and estimate EMG responses to unseen DBS frequencies. The model uses spectrograms as input features and this approach was chosen because neither the raw time series nor a Fourier frequency analysis alone could adequately capture the dataset’s temporal and frequency patterns [46]. Spectrograms, 2-D displays of frequency components over time windows, offer a balanced representation of temporal characteristics and frequency components [46]. Each value in a spectrogram is represented by a complex number, in the form of *a* + *jb*, where *a* is the real part and *b* is the imaginary part. The magnitude of this complex number, calculated as 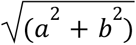 represents the strength or power of a particular frequency component at a specific time, indicating how dominant that frequency is in the signal during that time window. The phase, calculated as 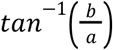, represents the phase angle of the frequency component, indicating the position of the wave relative to a time zero reference point. Thus, a spectrogram provides a time-resolved frequency analysis of a signal, showing both the magnitude and phase of each frequency component over time. Prior to constructing the spectrogram, the data underwent downsampling from 10,000 Hz to 1,000 Hz. This decision was informed by the power spectrum analysis and the Nyquist theorem, which indicated that the majority of the pertinent frequency information was concentrated below 500 Hz. Another characteristic displayed by the spectrogram is that the power in each frequency band doesn’t evolve over time, and so the frequency dynamics hold greater significance than the temporal dynamics. Therefore, spectrograms were constructed with each bin covering a frequency resolution of 1 Hz, and so with consideration to the Nyquist theorem, 501 frequency bins were created from a 1,000 Hz sampling rate. A 90% overlap was used to retain temporal information resulting in 101 time bins. The objective of the 1-D CNN model is to map the varying power levels and frequency patterns between the input Vim firing rate and output EMG for different DBS frequencies, therefore, the magnitudes were extracted from the complex spectrograms and processed whereas the phases were discarded. Figures 2 and 3 below illustrate the power spectrum for the Vim input and EMG output for each of the DBS frequencies.

**Figure 2.**
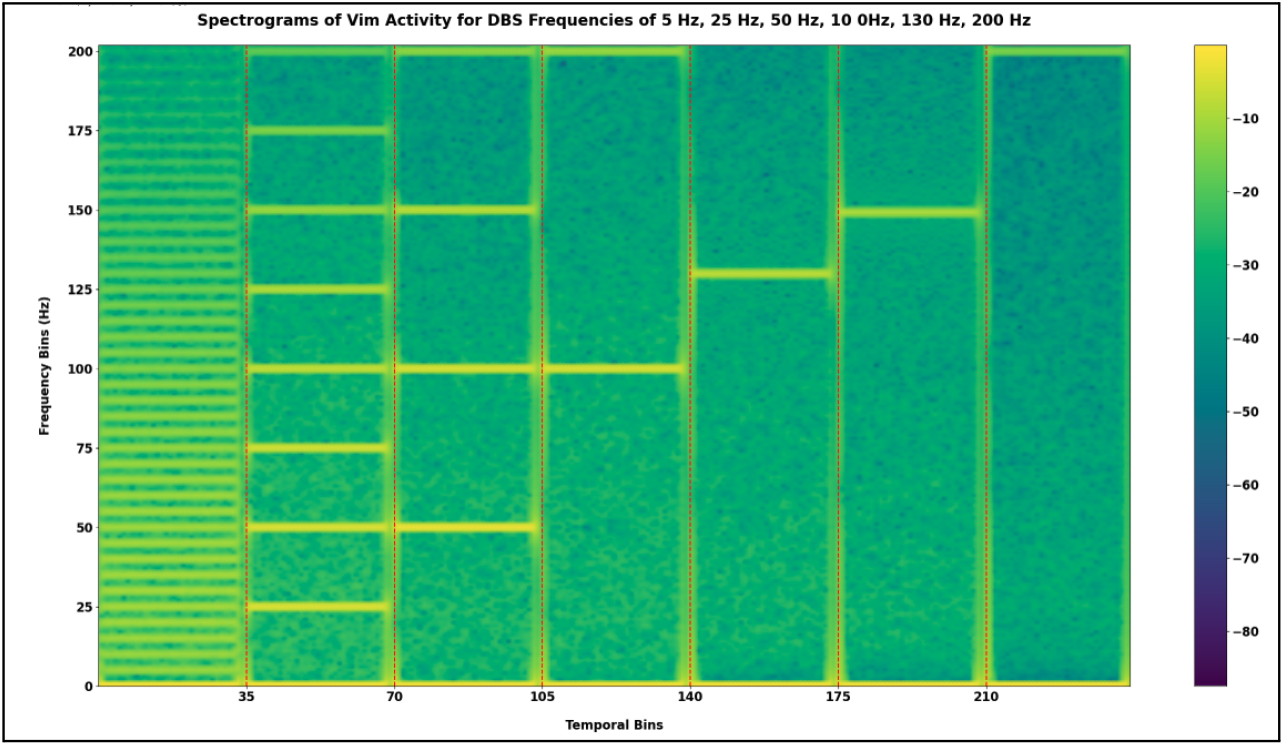
Concatenated spectrograms of **Vim** activity for DBS frequencies of 5 Hz, 25 Hz, 50 Hz, 100 Hz, 130 Hz, 200 Hz (separated by dotted red line).

**Figure 3.**
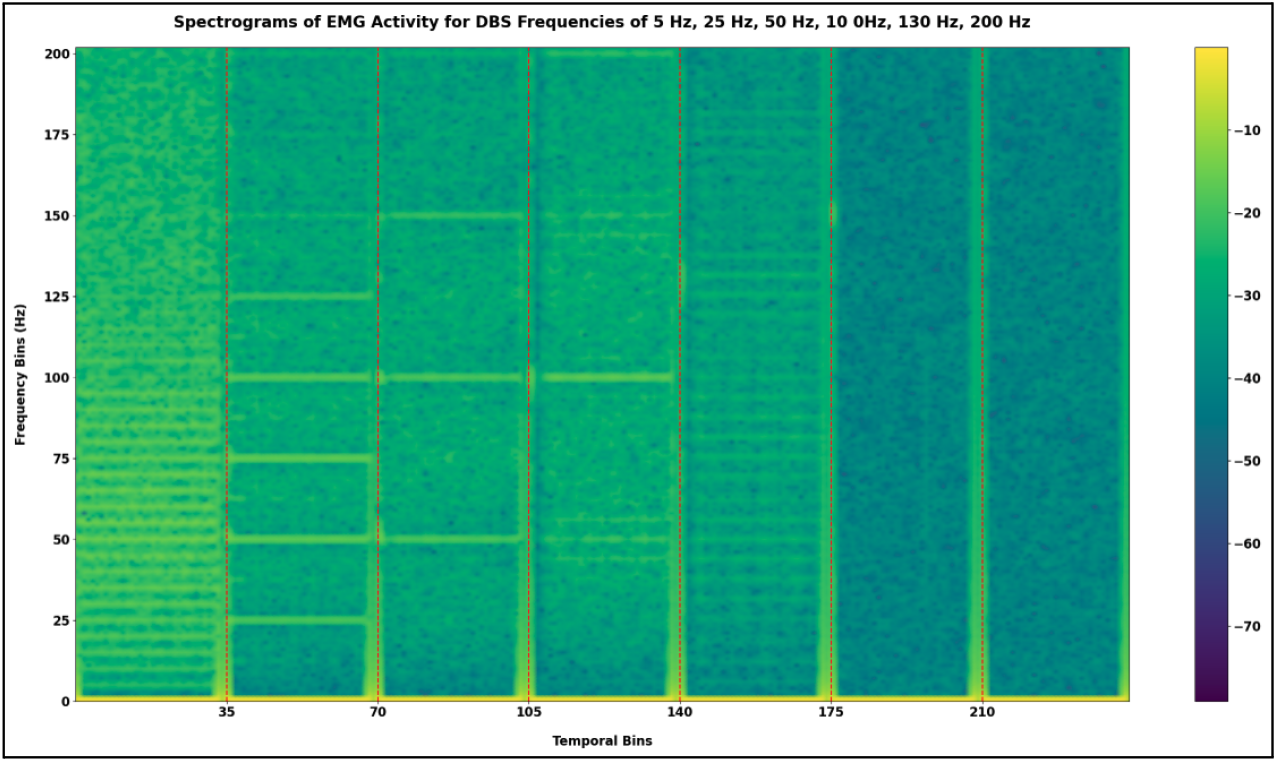
Concatenated spectrograms of **EMG** activity for DBS frequencies of 5 Hz, 25 Hz, 50 Hz, 100 Hz, 130 Hz, and 200 Hz (separated by dotted red line).

A clear distinction between the Vim and EMG data is the higher signal-to-noise ratio in the Vim activity across each of the DBS frequencies, as illustrated by the comparison of Figures 2 and 3. Another notable characteristic of the spectrogram data is that the majority of the power for both Vim and EMG is concentrated in the same frequency as the DBS frequency while exhibiting a decaying power trend for the harmonics of this frequency. However, DBS frequencies stimulated at 15 Hz and below have an outlier characteristic of the power being more evenly distributed across all the harmonics. Since these harmonics are more tightly spaced, the spectrograms for these low-frequency DBS resemble a random noise distribution as shown by the 5 Hz activity in the Vim and EMG in Figures 2 and 3. Consequently, incorporating these trials into the deep learning model training led to a decline in overall performance, likely due to the model mistaking noise for significant features. Therefore, these particular DBS frequency trials were discarded during the model training as frequencies below 20 Hz are not clinically effective anyway [31].

A 1-D CNN was used as the deep learning model for data-driven prediction because of its ability to capture local and global patterns across information in an image or in this case, a spectrogram [47]. A 1-D CNN was chosen over a 2-D CNN to reduce model complexity, as the spectrogram’s frequency content remained relatively stable over time [47]. Figure 1 displays the devised architecture of the convolutional network that was used. Each convolutional layer convolves the pre-existing layer with a kernel that outputs a learned feature map. The pooling layer reduces the dimensionality of the feature map by averaging across the time dimension. The output fully connected layer transforms the final feature map into an output predicted time series. The first layer in the architecture is a 1-D convolutional layer with 256 filters that process the input spectrogram features. Each filter is a set of learnable parameters that detect specific broad overall patterns in the spectrogram image along the frequency axis. Convolution is the process of applying these filters to the input data to produce feature maps, which highlight the presence of specific patterns detected by the filters. The kernel size, in this case, is 7, meaning each filter spans 7 frequency bins, enabling the detection of local patterns within that window. A ‘same’ padding is used to ensure that the output feature map has the same length as the input by adding zeros to the input boundaries. ReLU (Rectified Linear Unit) activation introduces non-linearity into the model by setting all negative values in the feature maps to zero, allowing the network to learn complex patterns faster and more efficiently (SOURCE). Following the first convolutional layer is a max pooling layer with a pool size of 2. Max pooling is a down-sampling technique that reduces the dimensionality of the feature maps by taking the maximum value within each window of the specified size (in this case, 2). This operation helps in reducing the computational load and controlling overfitting by retaining only the most prominent features. The output from this layer is then passed to a second 1-D convolutional layer with 512 filters, a kernel size of 3, ‘same’ padding, and ReLU activation. This layer functions similarly to the first convolutional layer but with a greater number of filters with a smaller kernel size, allowing the network to detect more complex finer frequency patterns. Both convolutional layers consist of a stride length of 1. Another max pooling layer with a pool size of 2 follows this convolutional layer, further reducing the data’s dimensionality. The resulting feature maps are then flattened into a one-dimensional vector, which is fed into a fully connected layer with 10,000 units and a linear activation function. This layer combines the features extracted by the convolutional layers to produce the final output of the network which is an output time series data of 10,000 points (a downsampled 10-second duration of EMG data). The fully connected layer intrinsically learns to perform an inverse transform on the spectrogram feature maps to output time series data. This architecture is designed to capture both local and global patterns in the frequency domain and how they evolve over time, making it suitable for analyzing both the temporal and spectral characteristics of the spectrogram.

A mean squared error was used as the loss function because of its efficacy in regression-based tasks with a learning rate of 0.0005 [48]. Finally, the output predicted EMG time series was compared with a test dataset from 20 to 200 Hz in intervals of 5 Hz. The R^2^ metric was used to evaluate the performance of the developed 1-D CNN model and also used to quantitatively compare against different deep learning models.

## Results

### EMG simulation from the encoding computational model

We simulated EMG from our encoding model, both during DBS-OFF and in response to Vim-DBS of different stimulation frequencies in [10, 200] Hz (**Figure 4**).

**Figure 4.**
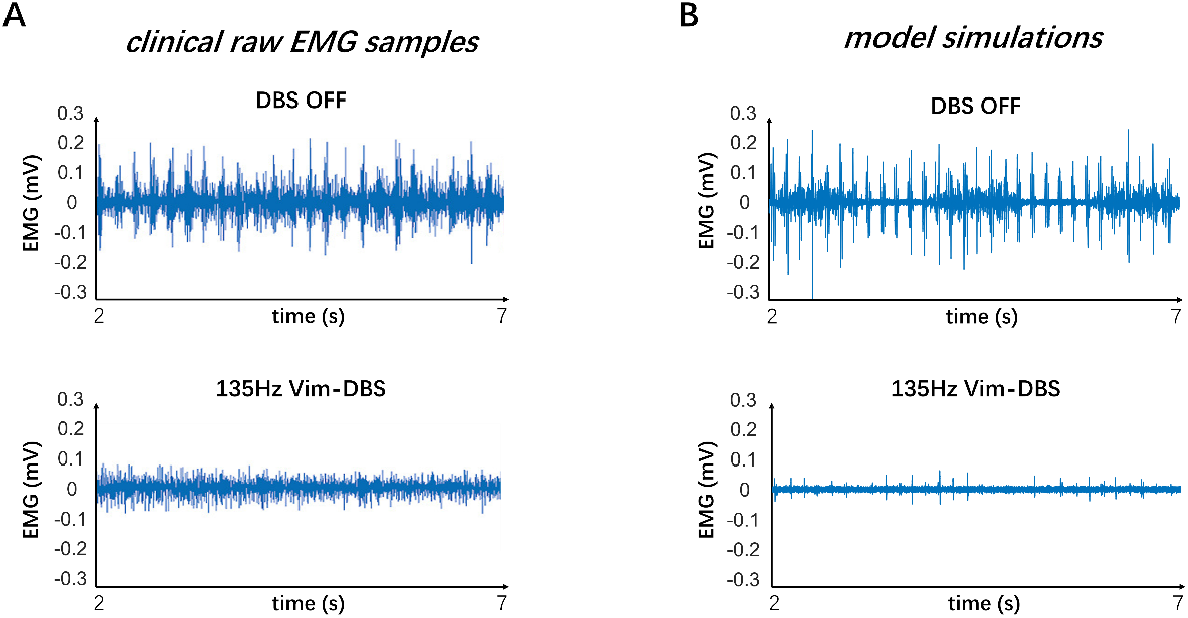
Compare clinical raw EMG samples and encoding model simulations. The clinical raw EMG samples were recorded from the extensor of an essential tremor patient during DBS-OFF and 135-Hz Vim-DBS (data from Cernera et al. (2021) [9]). We compare these clinical data with the steady-state EMG simulations from our encoding model.

The EMG simulation with DBS-OFF presented the typical tremor band (∼ 6 Hz) in the clinical EMG signals recorded from ET patients [33][34][18]. During DBS, in general, the EMG amplitude decreases as DBS frequency increases (**Figure 5**). The simulated EMG is mostly suppressed when DBS is delivered at a common clinical frequency ≥130 Hz [4][2][5] (**Figure 5**). We compared the model simulations with clinical data, which were raw EMG samples recorded from the extensor of an essential tremor patient during DBS-OFF and 135-Hz Vim-DBS (data from Cernera et al. (2021) [9]) (**Figure 4**). The model simulations were presented in steady-state beyond 2 seconds and were similar to these clinical data in terms of the EMG amplitudes (**Figure 4**). In clinical EMG, during DBS-OFF, we observed a 4 – 6 Hz tremor and a low voluntary muscle activity at ∼0.5 Hz; both observations are consistent with the model simulation (**Figure 4**). During 135-Hz Vim-DBS, in both clinical data and model simulation, we observed that the tremor activities are mostly suppressed (**Figure 4**). The tremor is less suppressed during 135-Hz Vim-DBS in clinical data compared with the model simulation (**Figure 4**). However, there is high variability in EMG activities across different individuals, and thus the clinical Vim-DBS frequency needs to be optimized specifically for individual ET patients [18][9][35].

**Figure 5.**
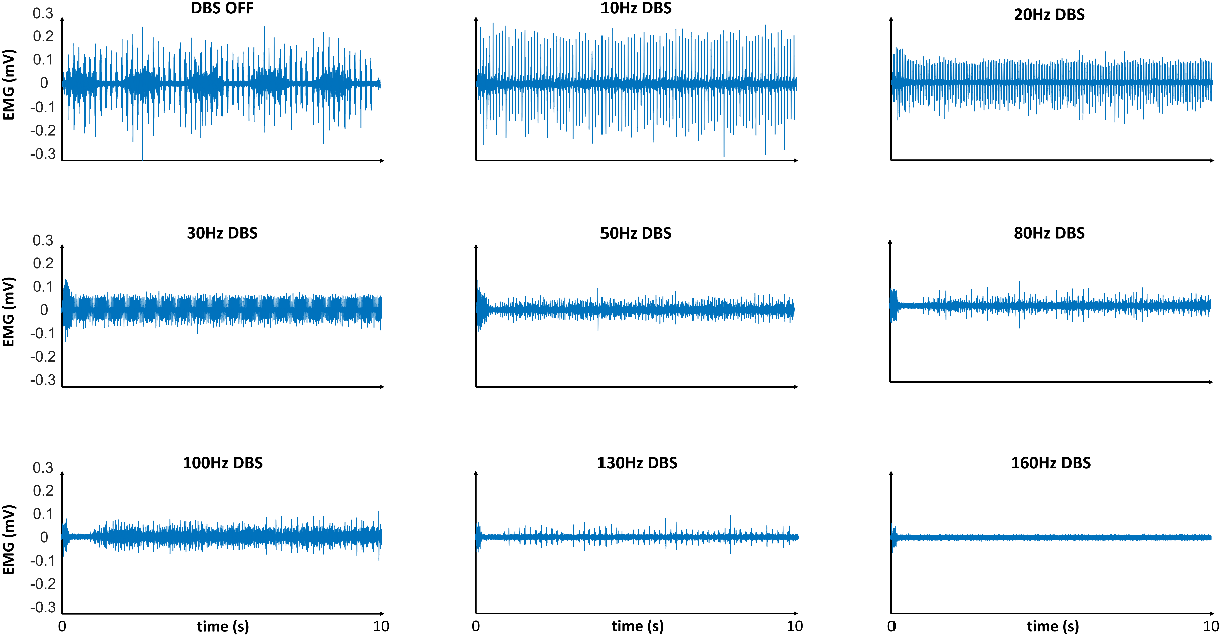
Model simulated EMG in response to different frequencies of DBS from our encoding model. These results are used for training a data-driven predictive deep learning model.

### EMG biomarker from the deep learning model

The EMG simulation from the encoding computational model is too slow for practical implementation. Thus, we developed the CNN deep learning model to predict the EMG biomarker from the encoding model, to facilitate the implementation speed. Testing results show that the data-driven CNN model fits the results from the encoding model very well. The EMG directly simulated from the encoding model is denoted as “reference EMG” (**Figure 5**). In **Figure 5**, we compared the CNN fit with the fit using a polynomial method, which was used in our previous work [16].

We observed that CNN fits the reference EMG well across the data from different DBS frequencies (**Figure 7**). We computed the *R*^2^ statistic [49] comparing the reference and fit in the frequency domain. We observed that in terms of the *R*^2^, the CNN fit (*R*^2^ = 0.997) is significantly more accurate than the polynomial fit (*R*^2^ = 0.277) (**Figure 7**). In particular, during low frequencies (≤ 30 Hz) of DBS, in terms of the bipolar symmetry feature of the EMG waveform, we observed that the CNN fit tracks this EMG feature more closely than the polynomial fit. Thus, the CNN deep learning model is much more accurate than the simple polynomial method in tracking the reference EMG and estimating EMG biomarkers for DBS frequencies between 20 and 200 Hz.

**Figure 7.**
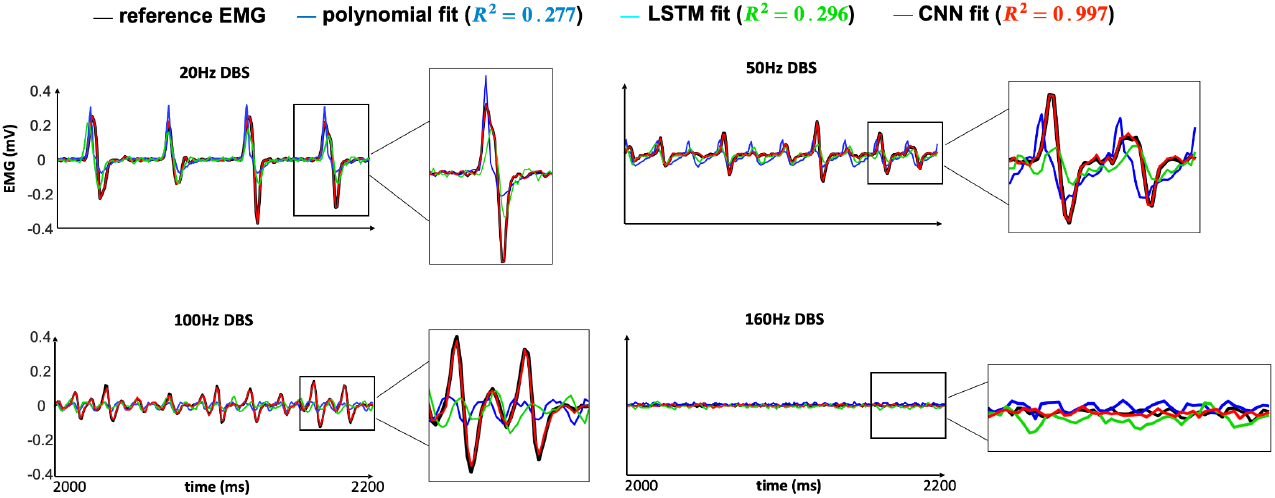
Comparison of the previously simulated EMG with the outputs generated from the polynomial fit model, LSTM model, and 1-D CNN model for various input DBS frequencies.

## Discussion

### Advantages of 1-D CNN using spectrogram features

Utilizing 1-D CNNs along the frequency axis of spectrograms presents several advantages, particularly in instances where the power within each frequency bin does not exhibit substantial temporal variation. One primary benefit is the ability of 1-D CNNs to efficiently capture and exploit local frequency patterns and features, which are crucial when the temporal dynamics are relatively static. By applying convolutions along the frequency axis, the model can discern fine-grained spectral characteristics and relationships across different frequency bands. This approach allows for the detection of subtle yet significant spectral features that may be indicative of underlying phenomena or conditions within the data. Moreover, the reduced complexity of 1-D convolutions compared to their 2-D counterparts leads to lower computational costs and faster training times, enhancing the feasibility of deploying these models in real-time or resource-constrained environments. Therefore, in cases where the temporal evolution of power is minimal, focusing on frequency resolution through 1-D CNNs along the frequency axis not only optimizes the model’s ability to learn relevant features but also provides a computationally efficient solution for spectral analysis.

### Other Deep Learning Approaches

As shown from the results by comparing R^2^ **(Figure 7)**, deep learning approaches show great potential for EMG waveform predictions compared to the previously developed polynomial fit model[16]. There is still an opportunity for improvement by optimizing the parameters during spectrogram computation and by tuning the deep learning model architecture. When choosing another deep learning approach for comparing performance with the 1-D CNN, the following methods were considered. Initially, the implementation of a deep learning model included a simple passing of the raw time series into a model which can capture the temporal dynamics of the data. RNNs are generally deep networks that are used to capture temporal dependencies in sequential data [50][51]. However, passing the data into an RNN network did not yield the best results. It seems that the long-term and short-term dependencies were not being properly captured, and so an LSTM network was implemented to address this issue. The bidirectional LSTM seemed to marginally increase the performance of the model and was therefore implemented in subsequent iterations. Although the bidirectional LSTM model showed improvement from the polynomial fit, qualitative analysis of the output demonstrated that it was still not capturing the relationship between Vim firing rate and EMG output very well in terms of both temporal characteristics as well as signal power when passing in the raw time series as features. Therefore, a Fourier spectrum of the Vim firing rate data was passed into the LSTM network. This iteration worsened the performance of the model and wasn’t capturing the signal frequency patterns effectively. In order to capture the Fourier components of the signal while maintaining temporal characteristics, spectrograms were constructed as features. This dramatically improved the performance of the model. Another model architectural component that was experimented with was identifying what aspect of this complex number to use as features for the aforementioned LSTM model training. Trials were conducted to identify the efficacy of using solely magnitude or concatenated features of the magnitude and phase, and the latter outperformed the others as features. The process was further improved when two independent LSTM blocks were used to independently process magnitude and phase, respectively, and then combined to produce an output complex spectrogram as shown by Figure 8.

**Figure 8.**
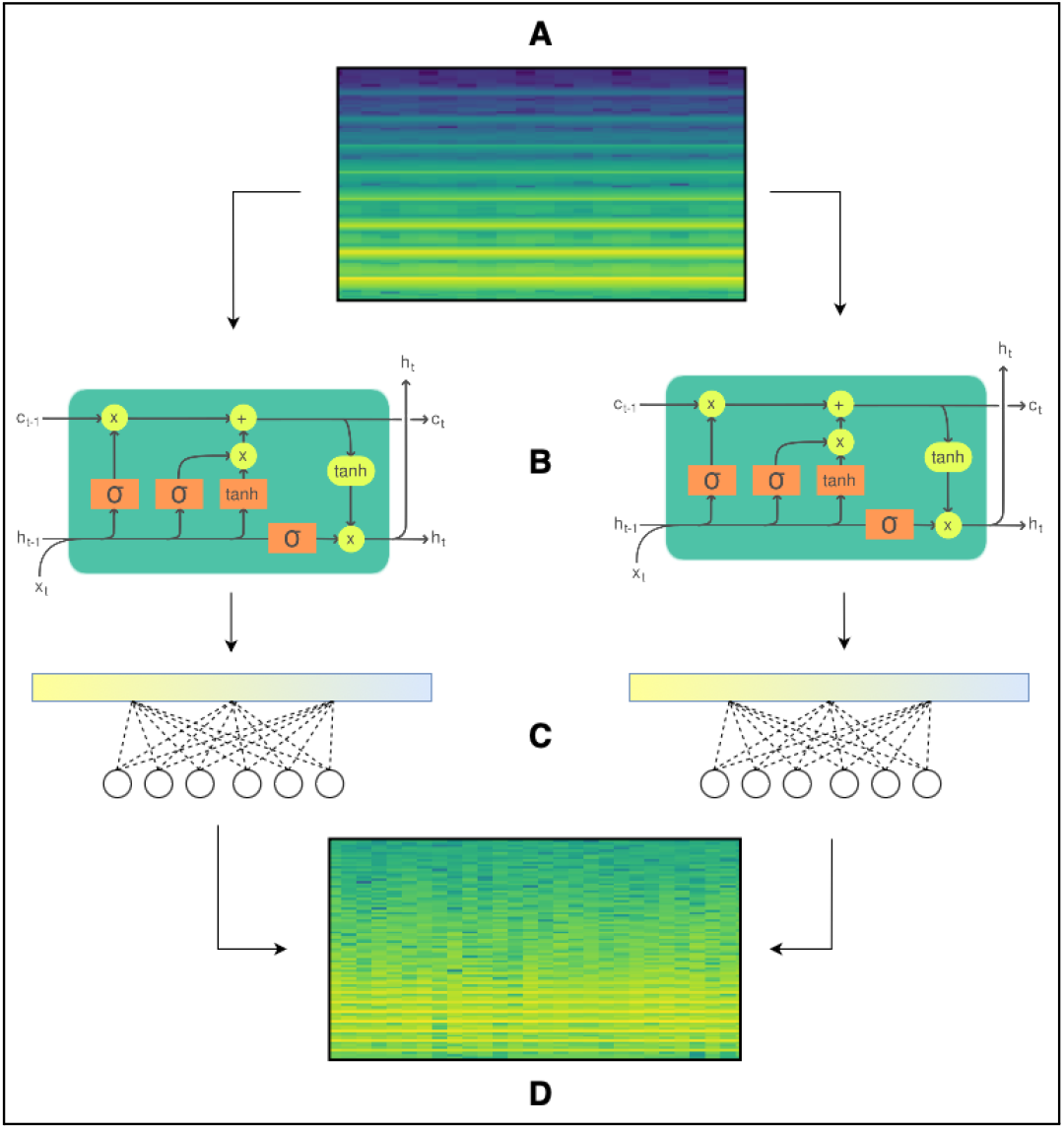
Architecture of the Bidirectional LSTM model that independently processes the Vim spectrogram to predict an EMG spectrogram. A) Input Vim Spectrograms for each DBS frequency where each value in the spectrogram is represented as a complex number. B) Two LSTM networks are used to process the magnitudes and phases of the input spectrograms separately. The LSTM consists of an input, forget, and output gate that collaboratively regulate the flow of information through the network’s cell state. The input gate controls which portions of the current input are stored in the cell, while the forget gate selectively erases irrelevant information from the previous cell state. The output gate determines what information is passed forward to the next hidden state, allowing the network to balance short- and long-term dependencies in the data. C) Fully connected layers processing the output of the LSTM models. D) Output predicted EMG spectrograms

Although the power levels for each of the DBS frequencies were somewhat captured, the frequency information itself was not sufficiently captured for each of the time series. As shown by the R^2^ performance, the 1-D CNN does a significantly better job at predicting the output time series compared to the bidirectional LSTM. The average R^2^ performance for the CNN model is 0.997 whereas the bidirectional LSTM model outputted an average R^2^ performance of 0.296.

Another deep learning method that might provide better results and worth investigating would be a transformer model [52]. This model would be able to input the entire time series data directly rather than a spectrogram which is a condensed version of the information, and it can capture both long-term and short-term dependencies which otherwise an LSTM model would not be able to retain. A limitation of transformer networks is that they require an abundance of data, but since the data used for model development and evaluation was simulated, it is possible to construct a sufficiently larger dataset. Another drawback of transformer models is their complexity as their architectures involve multiple layers of attention mechanisms and dense connections, which makes them challenging to interpret and less easily explainable [52]. Once a larger dataset has been assembled, it would be beneficial to compare transformers and the best-performing 1-D CNN network. Although deep learning is a ‘black box’ in nature, keeping the model architecture relatively simple addresses the issue of model explainability. In conclusion, deep learning shows potential for being a superior predictive model in comparison to the polynomial fit model because it can internally better understand the nonlinear relationship between Vim firing and EMG output. Furthermore, it is more accurate and flexible across a wider range of DBS frequencies.

### Improvements and Next Steps

To enhance the model’s robustness to different frequencies, especially those below 20 Hz, several strategies can be employed. One approach is to incorporate additional preprocessing steps that emphasize low-frequency components, such as applying bandpass filters to isolate and enhance these frequencies before generating the spectrogram. Additionally, augmenting the training data with variations that include low-frequency content can help the model learn to recognize these features more effectively. Improving the model’s architecture through experimentation with different convolutional layers and hyperparameters is also crucial. For instance, varying the kernel size, number of filters, and layer depth can help the model capture a broader range of spectral patterns. Employing techniques such as dropout and batch normalization can further enhance the model’s generalization capabilities and prevent overfitting. The construction of the spectrogram itself can be optimized to produce sharper and more informative representations. This can be achieved by adjusting parameters such as the window size, overlap, and type of window function used in the Short-Time Fourier Transform (STFT). A smaller window size might improve temporal resolution, while a larger window size might enhance frequency resolution, and so experimenting with this trade-off potentially may improve results. Experimenting with different window functions, such as Hamming or Hann windows, can also reduce spectral leakage and result in cleaner spectrograms. By fine-tuning these parameters and continually assessing the model’s performance, a more accurate and robust spectral analysis can be achieved which ultimately leads to improved model performance and reliability. The deep learning model was trained on simulated data, however clinically observed data is noisier, and so another key factor for evaluating the model’s robustness will be its ability to discern important features from noisier data. Therefore, artificially introducing noise into the spectrograms for training and then evaluating on clinically observed EMG spectrograms will be the next step in this data-driven predictive 1-D CNN deep learning model.

